# Is categorization in visual working memory a way to reduce mental effort? A pupillometry study

**DOI:** 10.1101/2021.11.23.469689

**Authors:** Cherie Zhou, Monicque M. Lorist, Sebastiaan Mathôt

## Abstract

Recent studies on visual working memory (VWM) have shown that visual information can be stored in VWM as continuous (e.g., a specific shade of red) as well as categorical representations (e.g., the general category red). It has been widely assumed, yet never directly tested, that continuous representations require more VWM mental effort than categorical representations; given limited VWM capacity, this would mean that fewer continuous, as compared to categorical, representations can be maintained simultaneously. We tested this assumption by measuring pupil size, as a proxy for mental effort, in a delayed estimation task. Participants memorized one to four ambiguous (boundaries between adjacent color categories) or prototypical colors to encourage continuous or categorical representations, respectively; after a delay, a probe indicated the location of the to-be-reported color. We found that, for set size 1, pupil size was larger while maintaining ambiguous as compared to prototypical colors, but without any difference in memory precision; this suggests that participants relied on an effortful continuous representation to maintain a single ambiguous color, thus resulting in pupil dilation while preserving precision. In contrast, for set size 2 and higher, pupil size was equally large while maintaining ambiguous and prototypical colors, but memory precision was now substantially reduced for ambiguous colors; this suggests that participants now also relied on categorical representations for ambiguous colors (which are by definition a poor fit to any category), thus reducing memory precision but not resulting in pupil dilation. Taken together, our results suggest that continuous representations are more effortful than categorical representations, and that very few continuous representations (perhaps only one) can be maintained simultaneously.

**Significance statement:** Visual working memory (VWM) can store visual information in two more-or-less distinct formats: continuous and categorical representations. It is widely assumed that VWM capacity for continuous representations is more limited than for categorical representations, yet this assumption has never been explicitly tested. Here we address this crucial question of whether continuous representations cost more resources, and as a consequence limits VWM capacity, as compared to categorical representations. To do so, we measured pupil size (as a proxy for mental effort) and memory performance in a delayed estimation task. Crucially, our results suggest that continuous representations are more effortful than categorical representations, and that only one continuous representation can be maintained in VWM at one time.

---

Visual information can be stored in visual working memory (VWM) as detailed, continuously varying representations, such as a specific shade of green, or as discrete, categorical representations, such as a prototypical green color. Traditionally, studies on VWM storage have focused on continuous representations. For example, one of the most commonly used analysis techniques in VWM research is the “mixture model” introduced by Zhang & Luck (2008). In this model, VWM performance is described as a mix of non-random responses that are centered on the memorized color with some random error (i.e. a von Mises distribution), and random responses (guesses) that are represented by a uniform distribution. Although this model provides a good way to quantify some characteristics of VWM storage, in its original form it does not take category boundaries or prototypes into account, and therefore implicitly assumes that all representations are continuous.

Recent studies have improved this by incorporating categorical representations into the mixture model (Bae & Luck, 2019; Hardman et al., 2017; Panichello et al., 2019; Pratte MS et al., 2017; Souza A.S. & Skora Z., 2017; Zhou et al., 2021). For example, Hardman and colleagues (2017) asked participants to memorize one or more colors, and to report the memorized color after a delay period. To analyze their results, the authors used a “revised mixture model”, which included a parameter that reflects the proportion of VWM items that are stored in continuous or categorical representations. Crucially, they found that only one continuous representation can be stored in VWM at a time, whereas multiple categorical representations can be stored simultaneously. In other words, their results suggest that VWM capacity for continuous representations is far more limited than for categorical representations.

The distinction between continuous and categorical representations is related to the emerging view of VWM as relying on different brain areas that are involved in different aspects of VWM maintenance (Christophel et al., 2017); specifically, continuous information would be represented predominantly in sensory areas, such as early visual cortex (Gayet et al., 2017; Hallenbeck et al., 2021; Harrison & Tong, 2009; but see Ester et al., 2015) whereas categorical information would be represented predominantly in non-sensory areas, such as parietal and prefrontal cortex, which are associated with learning and encoding categories at a more abstract level (Freedman et al., 2001; Lee et al., 2013; but see Ester et al., 2015). This involvement of multiple brain areas with diverse functions means that VWM can flexibly adopt different representational formats, such as continuous or categorical, to store visual information depending on the needs of the situation.

Although previous studies have shown *that* VWM can rely on both categorical and continuous representations, it is still an open question *why* VWM uses both types of representations. After all, if VWM is able to store a detailed representation, then why would people ever sacrifice this level of precision by resorting to a lower-resolution-categorical representation? Recent studies have started to address the question of why VWM switches between continuous and categorical representations. For example, in a study by Bae & Luck (2019), participants maintained a single orientation in VWM, while performing a visual discrimination task during the delay period. Crucially, the authors found that when VWM maintenance was interrupted by the discrimination task, memory responses became less precise and more biased towards the cardinal orientations (horizontal and vertical); that is, VWM representations became more categorical after a distracting task, possibly because the distraction consumed resources, thus reducing the resources available for VWM maintenance. This finding suggests that continuous representations, while more precise, are also more fragile than categorical representations.

In addition, as mentioned above, VWM representations are stored more categorically with increasing memory load (Hardman et al., 2017; Zhou et al., 2021). This suggests that VWM may have a lower capacity for continuous representations than for categorical representations; or, phrased differently, that it takes more resources to store a continuous representation than it does to store a categorical representation. (As we will get back to in the discussion, the first phrasing assumes a slot model of VWM, whereas the second phrasing assumes a resource model.) However, no study to date has directly tested this. Here we investigate this question by measuring pupil size while we encourage participants to maintain either continuous or categorical representations in VWM. Pupil dilation is commonly assumed to reflect mental effort during a range of cognitive tasks (Beatty & Lucero-Wagoner, 2000; Laeng et al., 2012; Mathôt, 2018; Unsworth & Robison, 2015), including working-memory tasks (Kahneman & Beatty, 1966). For example, a recent study found that when participants held one or more color items in VWM for a subsequent change detection task, their pupils became larger with increasing memory load (Unsworth & Robison, 2015). In the present study, we predict that maintaining continuous representations in VWM will consume more mental effort than maintaining categorical representations, and therefore that pupils will be larger while maintaining continuous information as compared to categorical information.

To test this assumption, we included two experiments: In Experiment 1, we identified prototypes for seven common color categories, as well as boundaries between adjacent categories, using a category assignment procedure. In Experiment 2, we asked participants to memorize one or more colors, and to reproduce one of the colors after a delay period. Crucially, the memory colors were either ‘ambiguous’ (i.e. the category boundaries, such as a hue that sits perfectly in-between red and orange) or prototypical (i.e. the category prototypes, such as a prototypical red hue), as identified in Experiment 1. We assumed that this manipulation would encourage the storage of continuous or categorical VWM representations, for respectively ambiguous and prototypical colors. Our main prediction was that pupil size should increase while maintaining ambiguous as compared to prototypical colors. Moreover, we also predicted a traditional effect of memory load; that is, pupil size should increase with increasing set sizes.

To foresee the results: our results suggest that participants were able to maintain a single continuous representation in VWM, even though this takes more mental effort than maintaining a single categorical representation; specifically, for set size 1, there was little difference in precision for ambiguous and prototypical colors, but there was a difference in pupil size, reflecting the additional mental effort required for maintaining continuous representations. However—strikingly—our results further suggest that participants failed to maintain multiple continuous representations, and instead resorted to categorical representations when memory load increased; specifically, for set sizes two and up, precision was lower for ambiguous as compared to prototypical colors, presumably because they were represented categorically despite not being a good fit to any category (see Starc et al., 2017); consistent with this interpretation, we now no longer observed a difference in pupil size, suggesting that participants used the same (or at least an equally effortful) representation for both ambiguous and prototypical colors. As we will discuss later, this is consistent with the notion that only one continuous representation can be maintained at a time, possibly because only a single item can be kept within the focus of attention in VWM (Olivers et al., 2011).

## Experiment 1

The purpose of Experiment 1 was to identify the prototype for each of the seven basic color categories (Hardman et al., 2017), as well as the boundaries between categories. On each trial, we showed one of 360 hues (every 1° of the hue-saturation-value color circle, with full saturation and value), and asked participants to assign them to one of seven color labels. The resulting prototypes and boundary values will be used as prototypical and ambiguous colors, respectively, in Experiment 2.

## Method

### Participants

Thirty-one first-year psychology students (aged from 18 to 26 years old; 26 female, 5 male) from the University of Groningen participated in this experiment online in exchange for course credits. All participants had normal or corrected-to-normal acuity and color vision. The study was approved by the Ethics Committee of Psychology of the University of Groningen (PSY-2021-S-0368).

### Procedure, data processing, and results

On each trial, a colored circle was shown in the center of the screen. The hues were chosen from a HSV (hue-saturation-value) color circle with full value (i.e., brightness) and saturation. Below the circle, seven color labels (red, pink, purple, blue, green, yellow, and orange) were shown horizontally (*Figure 1a*). The order of the color labels was random on every trial. Participants were instructed to click or tap on the label that they thought this hue belonged to. The circle and the color labels remained on the screen until response. Three-hundred-sixty different hues (for each 1° of the color circle) were shown once during the experiment in a random order.

**Figure 1.**
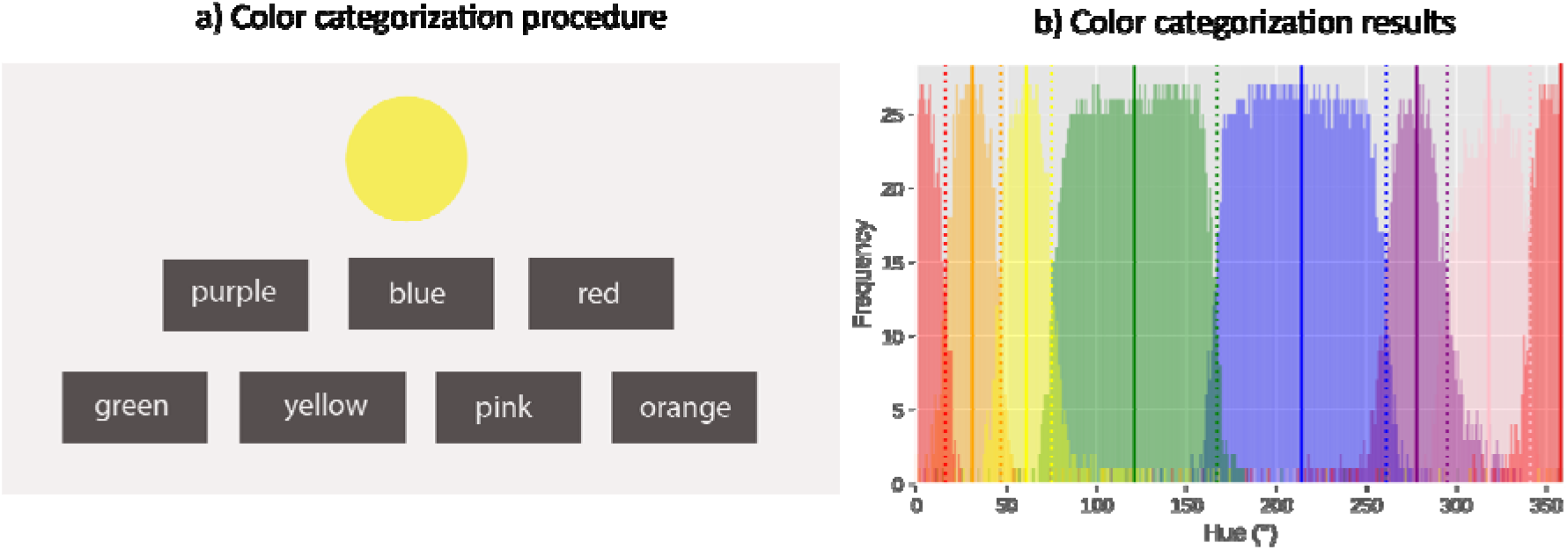
a) An example of the color categorization procedure in Experiment 1. b) Results of Experiment 1. Response frequency of each category is shown for each of the 360 hues. The center of each category (i.e. the prototypes) are marked by the vertical lines. The boundaries between categories are marked by the dotted lines. (Note: The category red is centered on 0° and is therefore split in this figure.)

Four participants whose responses strongly deviated from the average response (*Figure 1b*) were removed after visual inspection; the criterion for removal was subjective, but since removal was done before starting Experiment 2, this subjectivity did not correspond to ‘researchers degrees of freedom’ while conducting the main analyses. (See an example of excluded response distributions at https://osf.io/kjpey/.) Twenty-seven participants remained for further analysis.

Next, we calculated the response frequency of each category for every hue value, starting from red. When the most frequently assigned category changed from one hue value to the next, this was chosen as the boundary between the two categories. The average of two nearby boundary values was chosen as the prototypes of the category (*Figure 1b*; see details of response frequency of each category: https://osf.io/7sqne/).

## Experiment 2

## Method

### Participants

Thirty first-year psychology students (between 18 and 23 years old; 22 female, 8 male) from the University of Groningen participated in exchange for course credits. All participants had normal or corrected-to-normal acuity and color vision. The study was approved by the Ethics Committee of Psychology of the University of Groningen (PSY-2122-S-0014). Since we did not have an a-priori estimate of an expected effect size, our sample size was not based on a power analysis. Rather, we relied on our lab default of testing 30 participants for within-subject pupillometry studies. This is comparable to the rule of thumb recently provided by (Brysbaert & Stevens, 2018) of having 1,600 observations per condition (we have slightly more observations when considering our main effects, and slightly fewer when considering the full interaction).

### Stimuli, Design, and Procedure

Each trial started with a 1500 ms pre-cue consisting of a number of arrows indicating the locations of the stimuli that need to be memorized. (*Figure 2*). The number of arrows varied depending on the Memory Load condition (one to four). These arrows remained in the subsequently presented memory display, which showed four colors. (There were always four colors in the memory display, so that visual stimulation did not differ between different set sizes.) The memory colors were randomly drawn from a list of prototypical hue values from seven categories: red, pink, purple, blue, green, yellow, and orange; or from a list of seven ambiguous hue values that correspond to the boundaries between adjacent categories (e.g., red-pink, blue-green, etc.). The prototypes and the ambiguous colors are based on the values observed in Experiment 1. Next, following a 2500 ms retention interval, only one of the arrows from the previous displays remained on the screen for another 300 ms as a probe, indicating which color should be reproduced (i.e. target color). Finally, participants reproduced the probed memory color on a color circle with no time limit. Visual feedback followed, comparing what they had selected with the color that had been presented at the cued location.

**Figure 2.**
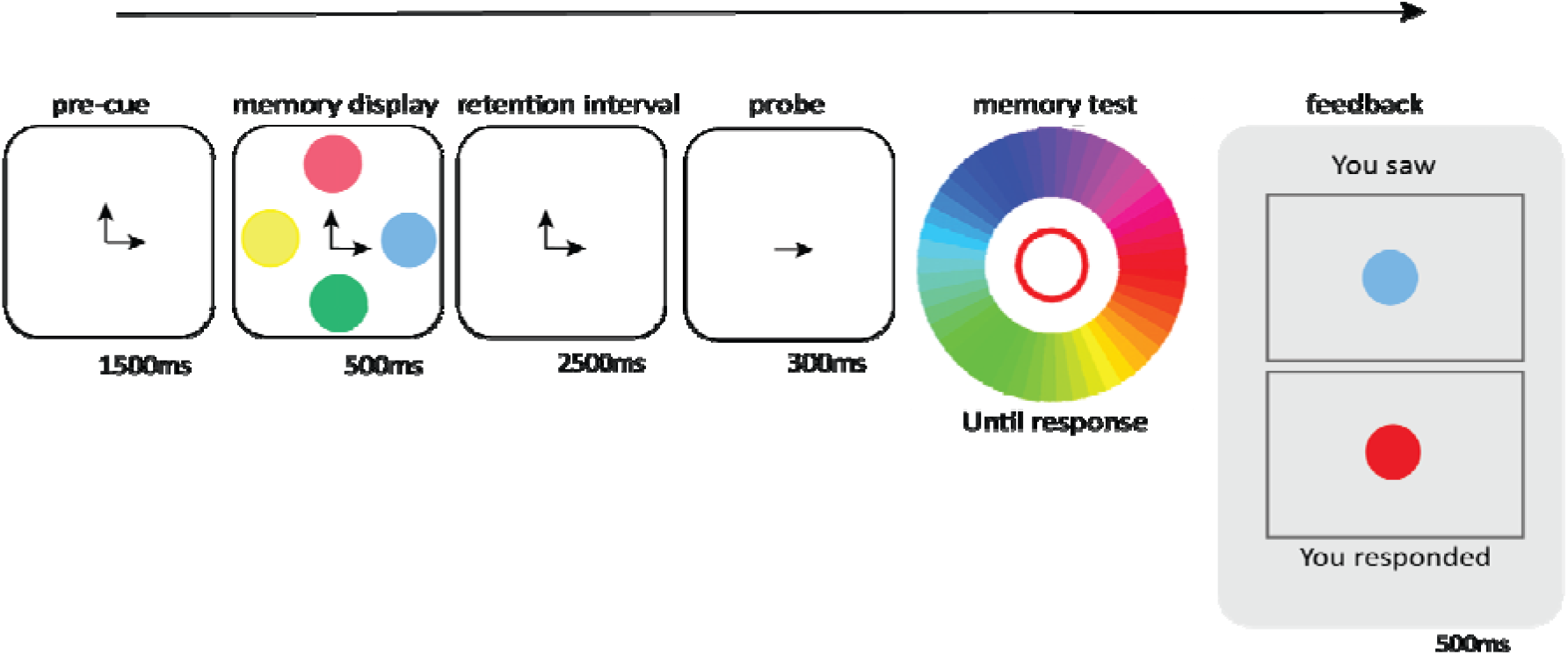
Sequence of events of a trial with a Memory Load of two, and a prototypical Color Type.

During the experiment, participants placed their chin on a head rest, at a distance of approximately 60 cm from the monitor. Pupil size was recorded monocularly with EyeLink 1000+ eye-tracker (SR Research, Ontario, Canada), with a sampling rate at 1,000 Hz. Gaze positions were calibrated using a nine-point calibration procedure. Before each trial, a drift correction was performed using a single-point recalibration. The experiment was conducted in a dimly lit room with no distractors. Stimulus presentation and response collection was controlled with OpenSesame (version 3.3.5; Mathôt et al., 2012) and PyGaze (Dalmaijer et al., 2014).

The Memory Load and Color Type conditions were mixed randomly within blocks without constraints. Participants completed 32 blocks of eight trials each (256 trials in total), preceded by one practice block of four trials.

### Preprocessing and exclusion criteria

First, blinks in the pupil data were corrected using cubic-spline interpolation (Mathôt, 2013). Next, pupil data was baseline-corrected by subtracting the mean pupil size during the presentation of the pre-cue from the entire pupil waveform. For each participant separately, we calculated the z-scored baseline pupil size; trials in which the baseline deviated more than 2 standard deviations from the mean were excluded from the pupil-size (but not the behavioral) analysis. In total, 348 trials were excluded based on this criterion. (The number of excluded trials per participant ranged from 8 to 17. See details here: https://osf.io/qvy6r/?pid=n9jgu.) Furthermore, we conducted a permutation test to test whether the mean accuracy of the participant was significantly above chance level. Specifically, we shuffled the response hue of all trials, and decided the “shuffled absolute error” based on the distance between the shuffled response hue value and the target hue value on each trial. We then conducted an independent samples t-test to test if the response error of each participant is significantly lower than the shuffled response error (with an alpha level of .05). No participants were excluded based on this criterion.

### Statistical analyses

#### Pupil Size

For each 100 ms window separately, we conducted a linear mixed effects model (LMER) using the R package lmerTest (Bates et al., 2018) with pupil size as dependent variable, Memory Load and Color Type as fixed effects, with random by-participant intercepts and slopes for all fixed effects. To qualify the Memory Load by Color Type interaction, we also conducted separate LMERs for each set size, with only Color Type as fixed effect. We used an alpha level of .05 for at least 200 ms (i.e. two consecutive windows).

#### Behavior: Precision and Guess Rate

First, we calculated the response error, which reflects the circular distance between the memory hue and the response hue. Next, we fitted a two-parameter mixture model, using the biased-memory toolkit (Zhou et al., 2021), to the distribution of response error, resulting in two parameters: Precision [0, 10000] and Guess Rate ([0, 1]). We conducted separate repeated measures analyses of variance (RM-ANOVA) with Memory Load and Color Type as independent variables, and each of the two model parameters as dependent variables: Precision and Guess Rate.

## Results

### Pupil Size

As shown in Figure 3a, there was an interaction between Color Type and Memory Load on pupil size (based on a criterion of *p* < .05 for at least a 200 ms contiguous interval as described above). This effect started at 1,300 ms and remained until 2,300 ms after the start of the retention interval. Specifically, at set size 1, pupils were larger when participants memorized ambiguous colors, compared to prototypical colors; at set sizes larger than 2, there was no difference in pupil size between the ambiguous and prototypical colors.

**Figure 3.**
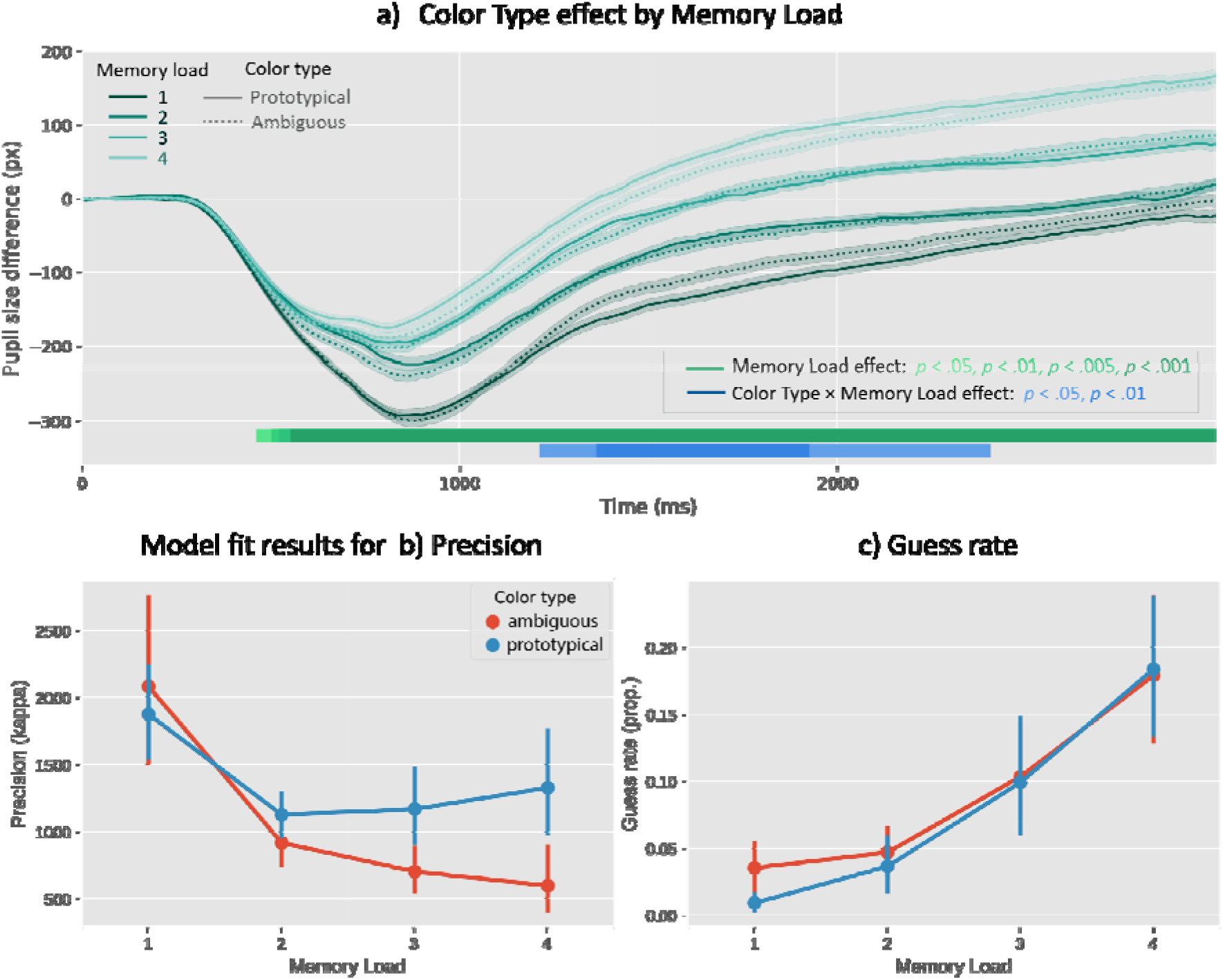
a) Pupil size since the onset of the memory display as a function of Color Type and Memory Load. b) Model fit results for Precision and c) Guess rate as a function of Color Type and Memory Load.

In addition, there was a strong Memory Load effect on pupil size that arose 500 ms after memory stimuli onset, and remained throughout the retention interval.

### Behavior: Precision and Guess Rate

For Precision, there was an effect of Color Type (*F*(1,29) = 5.21, *p* = .03), such that precision was higher for prototypical, as compared to ambiguous colors. Interestingly, we found an interaction effect between Color Type and Memory Load (*F*(3,87) = 3.17, *p* = .03; as shown in *Figure 3b*); specifically, at set size 1 and 2, there was no difference in Precision between ambiguous and prototypical conditions (1: *t*(29) = .58, *p* = .57; 2: *t*(29) *=* -2.02, *p* = .05); whereas at set size 3 and 4, responses in the ambiguous condition were significantly less precise than in the prototypical condition (3: *t*(29) = -2.83, *p* < .01; 4: *t*(29) = -3.14, *p* < .01). This suggests that when memory load was low, VWM relied on continuous representations for the ambiguous color; however, when memory load increased, VWM resorted to categorical representations even for the ambiguous color, which is reflected on the low precision for ambiguous colors. Moreover, there was a strong effect of Set Size on Precision (*F*(3,87) = 19.07, *p* < .01), such that precision decreased with increasing set size.

For Guess Rate, we found no effect of Color Type (*F*(1,29) = 1.08, *p* = .31), nor an interaction between Color Type and Memory Load (*F*(3,87) = .58, *p* = .63). However, we did find a strong effect of Set Size on Guess Rate (*F*(3,87) = 32.64, *p* < .01), such that guess rate increased with increasing set size (*Figure 3c*).

## Discussion

The present study investigated whether keeping continuous representations in VWM costs more mental effort than keeping categorical representations in VWM. To this end, we measured pupil size, as a proxy for mental effort, while participants maintained one or more prototypical or ambiguous colors in VWM; the prototypical colors corresponded to previously defined color-category prototypes, whereas ambiguous colors corresponded to the boundaries between adjacent color categories. (Color-category boundaries and prototypes had been determined in Experiment 1 using a separate group of participants.) The manipulation of prototypical versus ambiguous colors was designed to encourage categorical and continuous representations, respectively.

We found that, for set size 1, pupil size was larger in the ambiguous condition, as compared to the prototypical condition, presumably reflecting increased mental effort while maintaining a single continuous representation. For set size 1, there was no difference in memory precision between the two color-type conditions, suggesting that a continuous representation for a color that is difficult to categorize (e.g., shade that sits on the boundary between blue and green) has a comparable fidelity as a categorical representation for a color that is easy to categorize (e.g., a prototypical shade of blue). In contrast, for set size 2 and higher, there was no difference in pupil size between the ambiguous and prototypical conditions; however, precision for ambiguous colors was now substantially lower than that for prototypical colors, suggesting that VWM likely relied on categorical representations in all cases, also for the ambiguous colors that were difficult to categorize, thus resulting in reduced precision for these colors. In addition, we found a traditional effect of memory load on pupil size, reflecting that even for categorical representations, mental effort increases with memory load.

These results can be interpreted slightly differently following two opposing models of VWM: the slot model (Fukuda et al., 2010; Pratte MS et al., 2017; Zhang & Luck, 2008) and the resource model (Bays & Husain, 2008; Bays PM et al., 2009). According to the slot model, the capacity of VWM is limited to a fixed number of slots, with fewer slots for continuous representations, as compared to categorical representations (e.g., Hardman, 2017); as a result, when memory load reached the maximal number of slots for continuous representations (i.e. a single one in the case of the current study), VWM resorted to categorical representations. In contrast, the continuous resource model suggests that there is a single resource that is distributed across items; following this model, continuous representations would consume more resources than categorical representations. Although the slot-versus-resource debate is not our main focus in this study, our results are consistent with recent proposals (e.g., Pratte MS et al., 2017) that combine the characteristics of both models; that is, there are a limited number of slots for both continuous and categorical representations, and each slot has a variable precision.

Importantly, our findings are reminiscent of previous findings suggesting that only a single continuous representation can be maintained at a time (Hardman et al., 2017) and that, at least in some cases (Frătescu et al., 2019), only a single VWM representation can guide attention at a time (McElree, 2001; Oberauer, 2002; Olivers et al., 2011; van Moorselaar et al., 2014). In turn, these findings may be related to a recent proposal that VWM information can be maintained in two states: within or outside the focus of (internal) attention (Bae & Luck, 2019; Cowan, 2005; Oberauer, 2002). When there is only a single visual item to be maintained, this item can be maintained inside the focus of attention, which presumably corresponds to activity in visual cortex (Bae & Luck, 2019; Foster JJ et al., 2017; Harrison & Tong, 2009; Pasternak & Greenlee, 2005). Therefore, this item is stored as a rich, detailed representation in VWM. In turn, this corresponds to what we have referred to as a ‘continuous’ representation in this paper. The advantage of continuous representations is likely that they contain more sensory detail; however, maintaining continuous representations, as compared to categorical representations, comes at the expense of increased mental effort. Therefore, when more than one item is maintained in VWM, the additional items are maintained outside of the focus of attention, which presumably does not correspond to active processing in visual brain areas, but to activity in non-visual areas (e.g., prefrontal cortex; (Bae & Luck, 2019; Goldman-Rakic, 1995) or perhaps even to a type of ‘activity-silent’ representation (Stokes, 2015; Wolff et al., 2017). As a result, these additional items are stored as abstract, less-detailed representations in VWM. In turn, this corresponds to what we have referred to as ‘categorical’ representations. The advantage of such categorical representations is likely that they consume little mental effort; however, this comes at the expense of low precision, especially for items that are difficult to categorize (such as the ambiguous colors in the current study). Taken together, in recent years several theories of VWM have been proposed that are different from one another in some ways, yet all share the underlying notion that not all VWM representations are alike, and that the more detailed/ sensory/ continuous a VWM representation is: a) the more it is related to guidance of attention, b) the more it is related to activity in sensory brain areas; and c) as the present results suggest, the more mental effort it requires. An important avenue for future research and theorization will be to merge these loosely related notions into a coherent framework.

Overall, our results suggest that only a single continuous representation can be maintained in VWM at a time, and that maintaining a continuous representation costs more mental effort than maintaining a categorical representation; crucially, when multiple items need to be maintained, VWM resorts to categorical representations, which are less precise, especially for items that are not easily categorizable (such as our ambiguous colors), yet cost less mental effort.

### Open practices statement

All experimental data and materials can be found on the OSF (Open Science Framework): https://osf.io/n9jgu/. The Python implementation of the biased-memory toolbox can be found on GitHub: https://github.com/smathot/biased_memory_toolbox/.

## Reference

Bae, G., & Luck, S. J. (2019). What happens to an individual visual working memory representation when it is interrupted? British Journal of Psychology, 110(2), 268–287. https://doi.org/10.1111/bjop.12339

Bates, D., Kliegl, R., Vasishth, S., & Baayen, H. (2018). Parsimonious Mixed Models. 1506.04967 [Stat]. http://arxiv.org/abs/1506.04967

Bays, P. M., & Husain, M. (2008). Dynamic Shifts of Limited Working Memory Resources in Human Vision. Science, 321(5890), 851. https://doi.org/10.1126/science.1158023

Bays PM, Catalao RF, & Husain M. (2009). The precision of visual working memory is set by allocation of a shared resource. Journal of Vision, 9(10), 1–11. WorldCat.org. https://doi.org/10.1167/9.10.7

Beatty, J., & Lucero-Wagoner, B. (2000). The pupillary system. In Handbook of psychophysiology, 2nd ed. (pp. 142–162). Cambridge University Press.

Brysbaert, M., & Stevens, M. (2018). Power Analysis and Effect Size in Mixed Effects Models: A Tutorial. Journal of Cognition, 1(1), 9. https://doi.org/10.5334/joc.10

Christophel, T. B., Klink, P. C., Spitzer, B., Roelfsema, P. R., & Haynes, J.-D. (2017). The Distributed Nature of Working Memory. Trends in Cognitive Sciences, 21(2), 111–124. WorldCat.org.

Cowan, N. (2005). Working memory capacity. (pp. ix, 246). Psychology Press. https://doi.org/10.4324/9780203342398

Dalmaijer, E. S., Mathôt, S., & Van der Stigchel, S. (2014). PyGaze: An open-source, cross-platform toolbox for minimal-effort programming of eyetracking experiments. Behavior Research Methods, 46(4), 913–921. https://doi.org/10.3758/s13428-013-0422-2

Ester, E. F., Sprague, T. C., & Serences, J. T. (2015). Parietal and Frontal Cortex Encode Stimulus-Specific Mnemonic Representations during Visual Working Memory. Neuron, 87(4), 893–905. https://doi.org/10.1016/j.neuron.2015.07.013

Foster JJ, Bsales EM, Jaffe RJ, & Awh E. (2017). Alpha-Band Activity Reveals Spontaneous Representations of Spatial Position in Visual Working Memory. Current Biology□: CB, 27(20), 3216–3223. WorldCat.org. https://doi.org/10.1016/j.cub.2017.09.031

Frătescu, M., Van Moorselaar, D., & Mathôt, S. (2019). Can you have multiple attentional templates? Large-scale replications of Van Moorselaar, Theeuwes, and Olivers (2014) and Hollingworth and Beck (2016). Attention, Perception, & Psychophysics, 81(8), 2700–2709. https://doi.org/10.3758/s13414-019-01791-8

Freedman, D. J., Riesenhuber, M., Poggio, T., & Miller, E. K. (2001). Categorical Representation of Visual Stimuli in the Primate Prefrontal Cortex. Science, 291(5502), 312–316. https://doi.org/10.1126/science.291.5502.312

Fukuda, K., Awh, E., & Vogel, E. K. (2010). Discrete capacity limits in visual working memory. Current Opinion in Neurobiology, 20(2), 177–182. PubMed. https://doi.org/10.1016/j.conb.2010.03.005

Gayet, S., Guggenmos, M., Christophel, T. B., Haynes, J.-D., Paffen, C. L. E., Van der Stigchel, S., & Sterzer, P. (2017). Visual Working Memory Enhances the Neural Response to Matching Visual Input. The Journal of Neuroscience, 37(28), 6638. https://doi.org/10.1523/JNEUROSCI.3418-16.2017

Goldman-Rakic, P. S. (1995). Cellular basis of working memory. Neuron, 14(3), 477–485. https://doi.org/10.1016/0896-6273(95)90304-6

Hallenbeck, G. E., Sprague, T. C., Rahmati, M., Sreenivasan, K. K., & Curtis, C. E. (2021). Working memory representations in visual cortex mediate distraction effects. Nature Communications, 12(1), 4714. https://doi.org/10.1038/s41467-021-24973-1

Hardman, K. O., Vergauwe, E., & Ricker, T. J. (2017). Categorical working memory representations are used in delayed estimation of continuous colors. Journal of Experimental Psychology: Human Perception and Performance, 43(1), 30–54. https://doi.org/10.1037/xhp0000290

Harrison, S. A., & Tong, F. (2009). Decoding reveals the contents of visual working memory in early visual areas. Nature, 458(7238), 632–635. https://doi.org/10.1038/nature07832

Kahneman, D., & Beatty, J. (1966). Pupil Diameter and Load on Memory. Science, 154(3756), 1583. https://doi.org/10.1126/science.154.3756.1583

Laeng, B., Sirois, S., & Gredebäck, G. (2012). Pupillometry: A Window to the Preconscious? Perspectives on Psychological Science, 7(1), 18–27. https://doi.org/10.1177/1745691611427305

Lee, S.-H., Kravitz, D. J., & Baker, C. I. (2013). Goal-dependent dissociation of visual and prefrontal cortices during working memory. Nature Neuroscience, 16(8), 997–999. https://doi.org/10.1038/nn.3452

Mathôt, S. (2013). A simple way to reconstruct pupil size during eye blinks. Journal Contribution. https://doi.org/10.6084/m9.figshare.688001.v1

Mathôt, S. (2018). Pupillometry: Psychology, Physiology, and Function. Journal of Cognition, 1(1), 16. https://doi.org/10.5334/joc.18

Mathôt, S., Schreij, D., & Theeuwes, J. (2012). OpenSesame: An open-source, graphical experiment builder for the social sciences. Behavior Research Methods, 44(2), 314–324. https://doi.org/10.3758/s13428-011-0168-7

McElree, B. (2001). Working memory and focal attention. Journal of Experimental Psychology: Learning, Memory, and Cognition, 27(3), 817–835. https://doi.org/10.1037/0278-7393.27.3.817

Oberauer, K. (2002). Access to information in working memory: Exploring the focus of attention. Journal of Experimental Psychology: Learning, Memory, and Cognition, 28(3), 411–421. https://doi.org/10.1037/0278-7393.28.3.411

Olivers, C. N. L., Peters, J., Houtkamp, R., & Roelfsema, P. R. (2011). Different states in visual working memory: When it guides attention and when it does not. Trends in Cognitive Sciences, 15(7), 327–334. WorldCat.org. https://doi.org/10.1016/j.tics.2011.05.004

Panichello, M. F., DePasquale, B., Pillow, J. W., & Buschman, T. J. (2019). Error-correcting dynamics in visual working memory. Nature Communications, 10(1), 3366. https://doi.org/10.1038/s41467-019-11298-3

Pasternak, T., & Greenlee, M. W. (2005). Working memory in primate sensory systems. Nature Reviews Neuroscience, 6(2), 97–107. https://doi.org/10.1038/nrn1603

Pratte MS, Park YE, Rademaker RL, & Tong F. (2017). Accounting for stimulus-specific variation in precision reveals a discrete capacity limit in visual working memory. Journal of Experimental Psychology. Human Perception and Performance, 43(1), 6–17. WorldCat.org. https://doi.org/10.1037/xhp0000302

Souza A.S. & Skora Z. (2017). The interplay of language and visual perception in working memory. Cognition, 166, 277–297. WorldCat.org. https://doi.org/10.1016/j.cognition.2017.05.038

Starc, M., Anticevic, A., & Repovš, G. (2017). Fine-grained versus categorical: Pupil size differentiates between strategies for spatial working memory performance: Pupil size predicts working memory strategies. Psychophysiology, 54(5), 724–735. https://doi.org/10.1111/psyp.12828

Stokes, M. G. (2015). ‘Activity-silent’ working memory in prefrontal cortex: A dynamic coding framework. Trends in Cognitive Sciences, 19(7), 394–405. https://doi.org/10.1016/j.tics.2015.05.004

Unsworth, N., & Robison, M. K. (2015). Individual differences in the allocation of attention to items in working memory: Evidence from pupillometry. Psychonomic Bulletin & Review, 22(3), 757–765. https://doi.org/10.3758/s13423-014-0747-6

van Moorselaar, D., Theeuwes, J., & Olivers, C. N. L. (2014). In competition for the attentional template: Can multiple items within visual working memory guide attention? Journal of Experimental Psychology: Human Perception and Performance, 40(4), 1450–1464. https://doi.org/10.1037/a0036229

Wolff, M. J., Jochim, J., Akyürek, E. G., & Stokes, M. G. (2017). Dynamic hidden states underlying working-memory-guided behavior. Nature Neuroscience, 20(6), 864–871. PubMed. https://doi.org/10.1038/nn.4546

Zhang, W., & Luck, S. J. (2008). Discrete fixed-resolution representations in visual working memory. Nature, 453(7192), 233–235. https://doi.org/10.1038/nature06860

Zhou, C., Lorist, M. M., & Mathôt, S. (2021). Categorical bias in visual working memory: The effect of memory load and retention interval. Cortex. Registered report.

